# Numerical experiments for reconstructing cerebrospinal fluid flow based on contrast enhanced magnetic resonance images in the lower subarachnoid compartments of the brain

**DOI:** 10.1101/2025.04.10.648135

**Authors:** Martin Hornkjøl, Lars Magnus Valnes, Kent-André Mardal

## Abstract

In this paper we test various numerical methods for flow reconstruction based on both manufactured, idealized data and real patient data based on contrast enhanced magnetic resonance imaging after intrathecal contrast injection. As shown in previous studies, the imaging often display contrast gradients in localized regions although large areas have very small gradients. Velocities may as such be hard to assess in areas of low gradients. We compare optimal mass transfer with adjoint based data assimilation constrained by a convection diffusion equation. With well-chosen parameters the manufactured problems can be solved well in the idealized setting, but the performance is in general significantly worse in the patient specific setting. The methods predict maximal velocities well, but fail to reconstruct accurate velocity fields in areas without contrast gradients.

## 1 Introduction

The glymphatic system^1^ describes brain clearance facilitated by cerebrospinal fluid (CSF) in perivascular spaces (PVSs). Recently, it has become more evident that a crucial component of the system is the CSF flow in the subarachnoid space (SAS) and here the velocities are only partially known. In particular, the CSF flow in the SAS is complex and it pulsates at multiple frequencies with partially unknown magnitudes and directions. Directional and pulsating flows of the order of 20 *µm/s* in pial PVS at cardiac frequencies were measured in mice^2^. Furthermore, it has been demonstrated that the slow PVS pulsations during sleep was significantly larger in magnitude and resulting in similar or larger velocities^3^. This study was however not able to determine directionality. Using macroscopic magnetic resonance (MR) imaging, it has been estimated that the added dispersion effect of the combined pulsations are four orders of magnitude^4^, indicating that the CSF velocities within the SAS are a major components of the glymphatic system. Also here, the directionality was however not assessed.

In humans, the variation of CSF flows is perhaps even larger than in mice. Tracer transport of the order of three meter per hour^5,6^ have been estimate in the spinal SAS. However, less is know of the the velocities of the human cranial SAS. It has been shown, in patients with idiopathatic normal pressure hydrocephapuls (iNPH), that the contrast is mostly observed in the ventricle system^6^. However, the same study showed that contrast was often found within the cranial SAS rather than in the ventricle system on reference patients. The CSF flow within the ventricle system is rather well characterized^7–10^. It is known that the CSF flow in the aquaduct is of the orders of centimeter per second, and often significantly higher in iNPH patients. Furthermore, CSF flow in humans is heavily influenced by not only cardiac pulsations, but also respiration^8,11^. It is also clear that the CSF is modified by sleep and sleep-deprivation although the magnitude of change has not been quantified properly^12,13^. Finally, it has been shown that even a small directional bulk flow, as that of the chorioid plexus production, has the potential to speed up the clearance by orders of magnitude – from years to days (when combined with dispersion in the SAS)^14^. As such, it is interesting to investigate methodologies for assessing CSF flows numerically, based on human data.

Particle tracking velocimetry, a standard method in fluid dynamics, has been used to assess the flows^2^. This method has further been extended to physics informed neural networks^15^. Tracking dye or contrast is harder since only the gradients in the mixture can be tracked. For this reason, various alternative methods have been proposed to assess the velocity. For instance, optimal mass transfer (OMT) has been used in a series of papers^16–18^. The OMT methods are similar, although less general and potentially more efficient than the partial differential equation constrained optimization methods used in^19,20^ which requires optimization via adjoints. Hence, it is interesting to compare these different methods.

In this paper, we conduct a controlled study to evaluate several different methods for reconstructing velocities from images where contrast enhanced CSF spreads within the SAS of individuals undergoing so-called glymphatic magnetic resonance imaging (gMRI)^21,22^. The study will use manufactured solutions where the accuracy of the methods can be thoroughly assessed. Since the physical mechanisms of the glymphatic system on the macro-level in humans is only partially known, and in particular because it is unclear exactly how much the different production and absorption sites contribute in health and disease, we manufacture both seemingly plausible solutions as well as simplified solutions with less plausible physical foundations. We also, as detailed later, consider the early phases of contrast enhancement during gMRI as the flows are then mainly in the superior direction.

The methods we consider are partial differential equations (PDEs) based inverse modeling inspired by optical flows, both with and without restriction to the velocity field, and optimal mass transport methods, as proposed in several papers^23–26^. A main challenge with these methods is, however, that the movement of the contrast is just a surrogate of the underlying CSF flows. That is, CSF flow is only captured by the imaging by moving contrast gradients. Flow without contrast enhancement is as such invisible by the imaging, but may potentially be reconstructed by the constraining PDEs.

We find that while all methods recover velocities of the expected order of magnitude, none of the methods can accurately recover the complete velocity fields for all cases. The rest of the paper is laid out as follows. Section 2, defines the problems we want to solve and the methods we use to solve them. The results found with each method on each problem is presented in Section 3. Finally, the performance of the methods is discussed in Section 4 and the paper is concluded in Section 5.

## 2 Methods

We use two different optical flow methods, the optimal mass transport (OMT) method and three variations of the optimal control with diffusion and convention (OCDC) equations^27^ as constraints (Table 1). These methods will be employed to find the velocity field in; a box mesh with manufactured solutions, a realistic mesh with manufactured solutions and a realistic mesh with specific acquired data. Of the three OCDC methods used, two of them has their velocity field restricted. One restricted to be purly in the z-direction (OCDCz method) and one following a Stokes field (OCDCs method). A main motivation behind these methods is to simplify the minimization, assuming we have *a priori* knowledge of the flow, such as its main direction, the OCDCz method, or that it is flow of Stokes type with known inflow and efflux locations. The methods will be defined in detail below.

**Table 1.** The different methods used in this paper. The methods are defined in terms of their baseline method, the constraints on the velocity field and whether or not they have a regularization term. In total, there are three OCDC type methods and one OMT type method.

| Methods | Baseline method | Velocity field constraint | Regularization |
| --- | --- | --- | --- |
| OCDC | OCDC | None | True |
| OCDC<sub>z</sub> | OCDC | $\phi_z$ | True |
| OCDC<sub>s</sub> | OCDC | $\phi_s$ | False |
| OMT | OMT | None | True |

The optimization is based on tracer concentration images, both from manufactured problems and patient data. The goal with the manufactured problem is to test the validity of the different optical flow methods in a controlled environment. The manufactured problems are solved on a simple 2 cm x 1 cm x 1 cm box to look at performance in a simplified optimal case, and on a realistic patient mesh. We look at manufactured problems constructed with different types of velocity fields, and we also look at problems with added noise. In total, we construct seven different problems; The box mesh with velocity field in the z-direction (BZ) problem, the box mesh with a Stokes velocity field (BS) problem, the box mesh with velocity field in the z-direction and added noise (NBZ) problem, the patient mesh with velocity field in the z-direction (PZ) problem, the patient mesh with a Stokes velocity field (PS) problem, the patient mesh with velocity field in the z-direction and added noise (NPZ) problems, and the patient specific data (PSD) problem (Table 2). All these problems will be defined in the following sections.

**Table 2.** The different problems explored throughout the paper. Each problem is defined by the mesh used, the velocity field used when creating the concentration images and whether the problem has added noise. There are three manufactured problems on the box mesh and three for the patient mesh. In addition, there is the PSD problem on the patient mesh.

| Problems | Mesh | True velocity field | Noise |
| --- | --- | --- | --- |
| BZ | Box | $\phi_z$ | False |
| BS | Box | $\phi_s$ | False |
| NBZ | Box | $\phi_z$ | True |
| PZ | Patient | $\phi_z$ | False |
| PS | Patient | $\phi_s$ | False |
| NPZ | Patient | $\phi_z$ | True |
| PSD | Patient | Unknown | False |

In total, we test four different optimal flow methods on seven different problems (three manufactured problems on a box mesh, three manufactured problems on a patient mesh and one based on actual patient data). This is done for several values of mesh resolutions, regularization parameters and timestep lengths between the concentration images.

### 2.1 Box mesh

The box meshes have dimensions 2 cm x 1 cm x 1 cm. We create three box meshes with resolutions 6144, 12000 and 26364 cells. The bottom, top and sides of the mesh (Figure 1**A**) are marked for future calculations. Note that in the method description to follow Ω_inside_ is zero for the box case.

**Figure 1.**
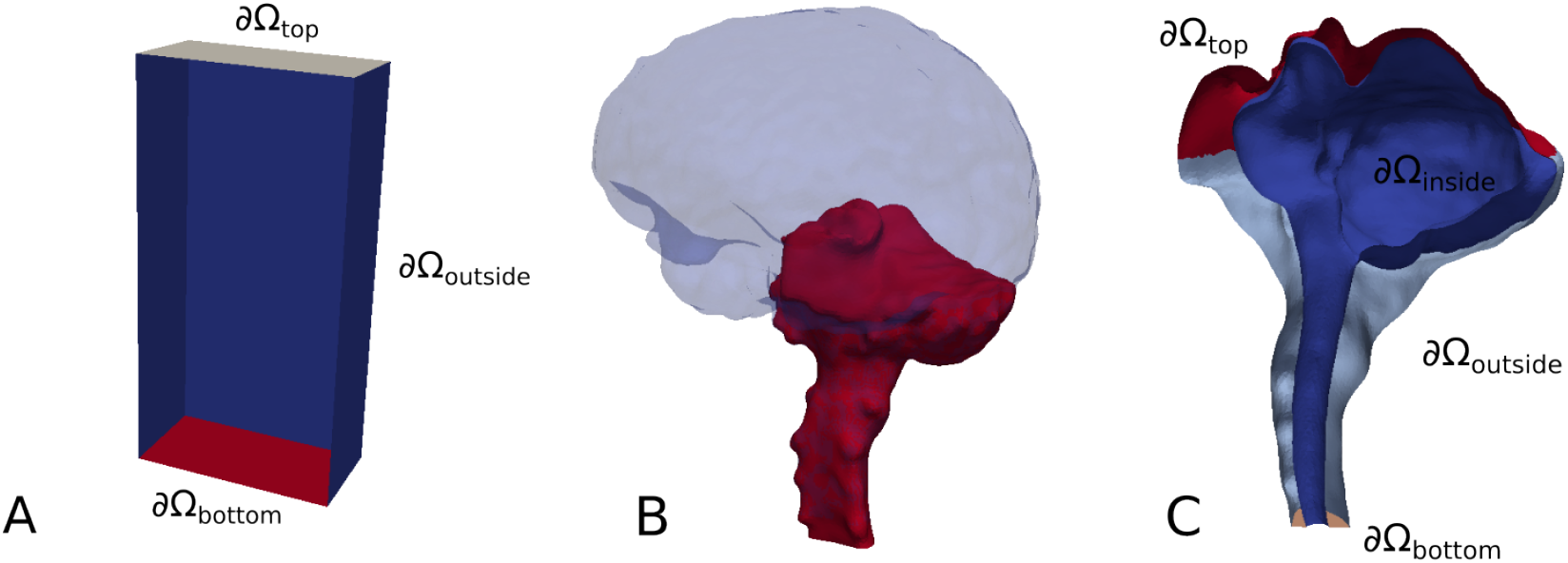
(**A**) The boundaries used for calculations in Section 2 for the box mesh. ∂ Ω_bottom_ is marked in red, ∂ Ω_outside_ in blue, and ∂ Ω_top_ in grey. (**B**) The part of the intracranial compartment that is the focus of this paper is marked in red, in opposition the full of the intracranial compartment marked in blue. (**C**) The boundaries used for calculations in Section 2. ∂ Ω_top_ is marked in red, ∂ Ω_inside_ in blue, ∂ Ω_outside_ in light blue and ∂ Ω_bottom_ in orange.

### 2.2 Patient data and meshes

We use baseline T1-weighted (T1w) and T2-weighted (T2w) MR images of a patient from a previous study. Starting from these T1w and T2w images of the patient, we use FreeSurfer to extract surfaces and choose to only look at the brain stem and cerebellum region (Figure 1**B**). We are only interested in the SAS space, so we exclude the brain stem and cerebellum. The surfaces has several holes, which we manually repair by smoothing the area around it. In order to have a flat bottom in the brain stem, we cut the mesh 5 mm above its lowest point. Additionally, the SAS is somewhat expanded, mostly between 1 and 2.5 mm depending on the area. This results in a thin zero layer on the edge of the mesh after interpolation of the images. When setting boundary conditions, we map the values on the images that are closest to the boundary onto the boundary. For the patient, three mesh resolutions are created with roughly 1, 2 and 4 million cells.

For the patient data, used in the following sections, we use two images at times 09:56:20 and 10:54:08, which equates to a time difference of 0:57:48.

### 2.3 Manufactured problems

In order to test our methods in a controlled environment, we solve the convection-diffusion equation to create concentration images for a known velocity field. We create two manufactured problems for both the box mesh and the patient mesh. One where the velocity field is set to *ϕ* (*x*) = *ϕ*_*z*_ = (0, 0, 20) mm/h, to be referred to as the BZ and PZ problem for the box mesh and the patient mesh respectively, and one where *ϕ* (*x*) = *ϕ*_*s*_(*x*) is the solution of the stokes equation, to be described in Section 2.5. The latter will be called the BS problem for the box mesh and PS problem for the patient mesh. The convection-diffusion equation used is as follows

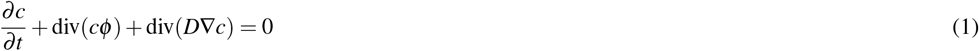

with boundary and initial conditions conditions

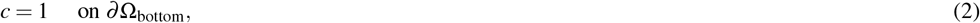

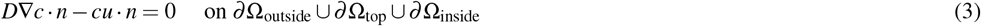

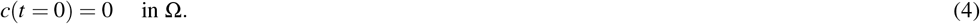

where Ω_bottom_ and Ω_outside_ are the bottom, sides and top of the box mesh. Ω_inside_ is zero for the box mesh. For the patient mesh Ω_bottom_ is the bottom surface of the spine, Ω_inside_ is the boundary facing the brain parenchyma, Ω_outside_ is the boundary facing the skull and Ω_top_ is, roughly, the boundary facing the rest of the intracranial compartment (Figure 1**C**). *D* = 3.8 · 10^−4^ mm^2^/s is the free water diffusion coefficient^19^. The choice *c* = 1 is arbitrary and the manufactured solution is scaled such that the total mass is the same at the final time as in the second data image. The convection-diffusion equation is solved using the finite element method (FEM).

The simulation time is 45 minutes and 6 hours for the box and patient mesh, respectively. The OMT and OCDC methods, described in Sections 2.4 and 2.6, is based on two scalar fields *I*_0_(*x*) and *I*_1_(*x*) representing concentrations *c*_0_ and *c*_1_ at times *t*_0_ and *t*_1_. We choose *t*_0_ to be 22.5 minutes for the box mesh and 5 hours for the patient mesh and consider a range of *t*_1_ choices.

By inspection of representative regions in the patient specific data, we observe a change of *±* 30-40 % in concentration. We want to test how this level of noise affects our methods when recreating the manufactured solutions. This is done by changing the concentration of our manufactured solution by up to *±* 35 % of the maximum concentration at all points, uniformly distributed. This is only done for the manufactured solution with velocity field *ϕ* (*x*) = (0, 0, 20) mm/h. The problem with added noise is referred to as the NBZ and NPZ problem for the box mesh and patient mesh respectively.

### 2.4 The OCDC method

Let *I*_0_(*x*) and *I*_1_(*x*) be two scalar fields representing two concentrations, *c*_0_ and *c*_1_, at two different times *t*_0_ and *t*_1_ respectively. We denote the time difference as Δ*t* = *t*_1_ − *t*_0_. The velocity field moving the concentration from *c*_0_ at *t*_0_ to *c*_1_ at *t*_1_ is called *ϕ*.

We assume that the velocity field *ϕ* (*x,t*) and concentration *c*(*x,t*) satisfy the convection-diffusion equation

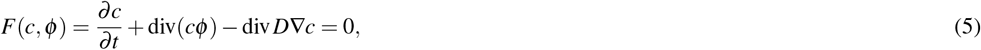

with the boundary and initial conditions

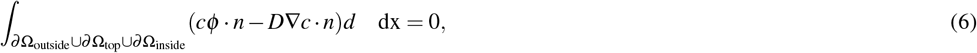

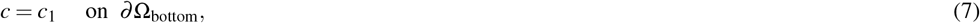

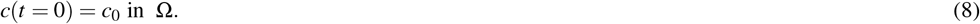

Solving *F* for *c* given a *ϕ* will be referred to as solving the forward problem.

Given the concentration observations *c*_0_ and *c*_1_, we want to find the *c* and *ϕ* that minimizes the objective functional

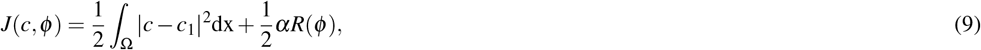

where the initial value for *c* is *c*_0_ and the initial guess for *ϕ* is 0, while still satisfying (5). *R* is a regularization term and *α* is a weight that determines the regularization strength. We choose *R* to be

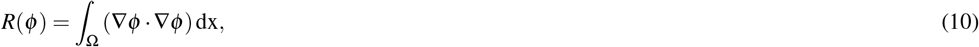

which minimized the gradient of the velocity field. The value of the functional *J* is the difference between the found and the true solution.

We can modify the OCDC method by adding restrictions to the velocity field *ϕ*. Using this, we introduce a variant of the OCDC method, called the OCDCz method, where *ϕ* = *ϕ*_*z*_ = *β* (*x*)*i*_*z*_ and optimize with respect to the scalar function *β* (*x*). That is, we force the velocity field to be strictly in the *z*-direction. As we expect the flow in this region of the brain to adhere to the Stokes equation, we also create the OCDCs method where *ϕ* = *ϕ*_*s*_ = *Cu*. Here, *u* is the solution of the Stokes equation, as described below in Section 2.5, and we optimize with respect to the scalar *C*. For the OCDCs method the regularization parameter is zero.

### 2.5 Stokes velocity field

Earlier in this section, we defined the BS and PS problem and the OCDCs method. These all rely on a velocity field created by solving the Stokes equation

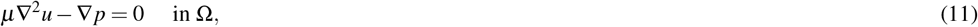

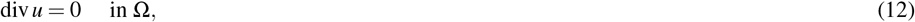

where *µ* = 8 *×* 10^−4^ is the viscosity. We choose boundary conditions

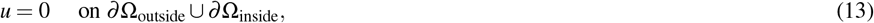

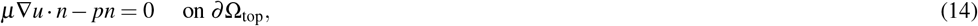

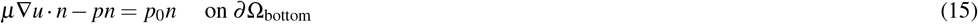

and solve the set of equations with the FEM. The choice of *p*_0_ is arbitrary, but the solution is scaled so that the maximum velocity for *u* is 20 mm/h.

### 2.6 OMT with kinetic energy minimization

Another method tested is the OMT method with kinetic energy minimization. We assume that the concentration transport by the velocity field *ϕ* is governed by the continuity equation

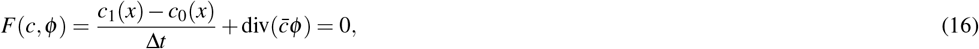

following^23^, where Δ*t* is the time between the images *I*_0_ and *I*_1_, which define the concentrations *c*_0_ and *c*_1_, and 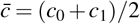 is the average concentration. We also define a regularization which minimize the kinetic energy of *ϕ* as

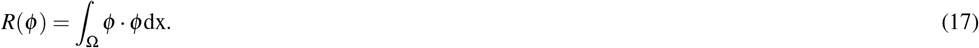

Note that the regularization does not include the concentration, since some regions of our domain have zero concentration. With these equations, we define a minimization problem

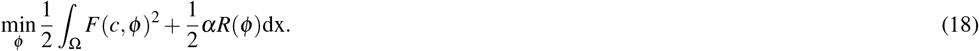

This can be rewritten to a variational formulation

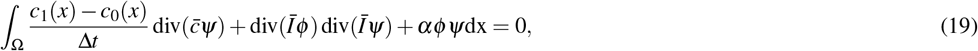

following^27^, which is solved for the unknown *ϕ*.

### 2.7 *L*_2_ difference

In the coming sections, we will use a normalized *L*_2_ difference to determine the correctness of velocity fields and the error caused by noise. By the normalized *L*_2_ difference we here mean

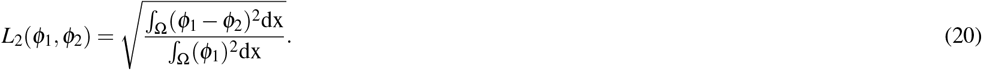

where *ϕ*_1_ and *ϕ*_2_ are either scalar or vector fields.

### 2.8 Summary of Problems, Methods and Parameters

In total, we have looked at two different types of meshes; a box mesh and a realistic mesh. For the box mesh, we are trying to recover the velocity fields for three different manufactured problems (Figure 2) and for the patient mesh, we are looking at the same manufactured problems (Figure 3**A**) in addition to the patient data problem (Figure 3**B**). The first manufactured problem is based on a velocity field in *z*-direction, the second a manufactured problem is based on solving the Stokes equation and the third manufactured problem is based on a velocity field in the *z*-direction with a uniform noise of *±* 35 % of the max concentration at all points(Table 2). The methods employed are OCDC, OCDCz, OCDCs and OMT methods (Table 1). Each of the methods depend on two parameters (except for the OCDCs method where the regularization is always zero), namely, the regularization parameter *α* and the time between the two images used for the inverse method. For the manufactured problem we look at different timesteps Δ*t* = [5.67, 11.25, 16.92, 22.5] minutes for the box mesh and Δ*t* = [15, 30, 45, 60] minutes for the patient mesh. For the PSD problem the timestep is always Δ*t* = 0:57:48. The regularization parameter is chosen to be *α* = [10^−4^, 10^−5^, 10^−6^] for BZ, BS and NBZ problems, *α* = [10^−6^, 10^−7^, 10^−8^] for the PZ, PS and NPZ problems and *α* = [10^−4^, 10^−5^, 10^−6^] for the PSD problem. An exception is the OCDCs method where there is no regularization as the change in gradient is predetermined by the Stokes field. The choices of *α* are based on numerical experimentation. Each problem has also been solved for 3 different mesh resolutions of 6144, 12000 and 26364 cells for the box and roughly 1, 2 and 4 million cells for the patient mesh.

**Figure 2.**
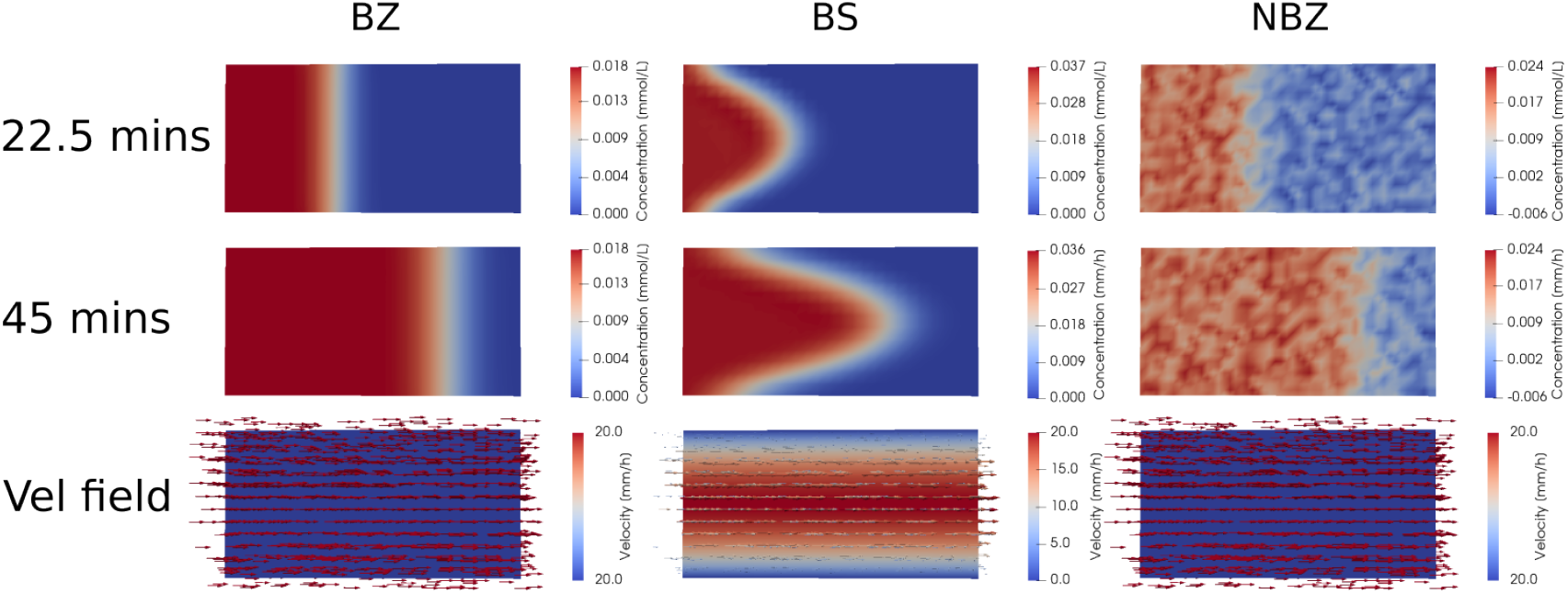
The manufactured concentration images used for the BS, BZ and NBZ problems (top two rows) with the corresponding velocity fields used to create them (bottom row). The second image at 45 minutes is the longest computed timestep. The boxes are cut in half, such that we are seeing the center of the box. Ω_bottom_ is to the left and Ω_top_ is to the right. Note that the velocity field for the BZ and NBZ problem is the same.

**Figure 3.**
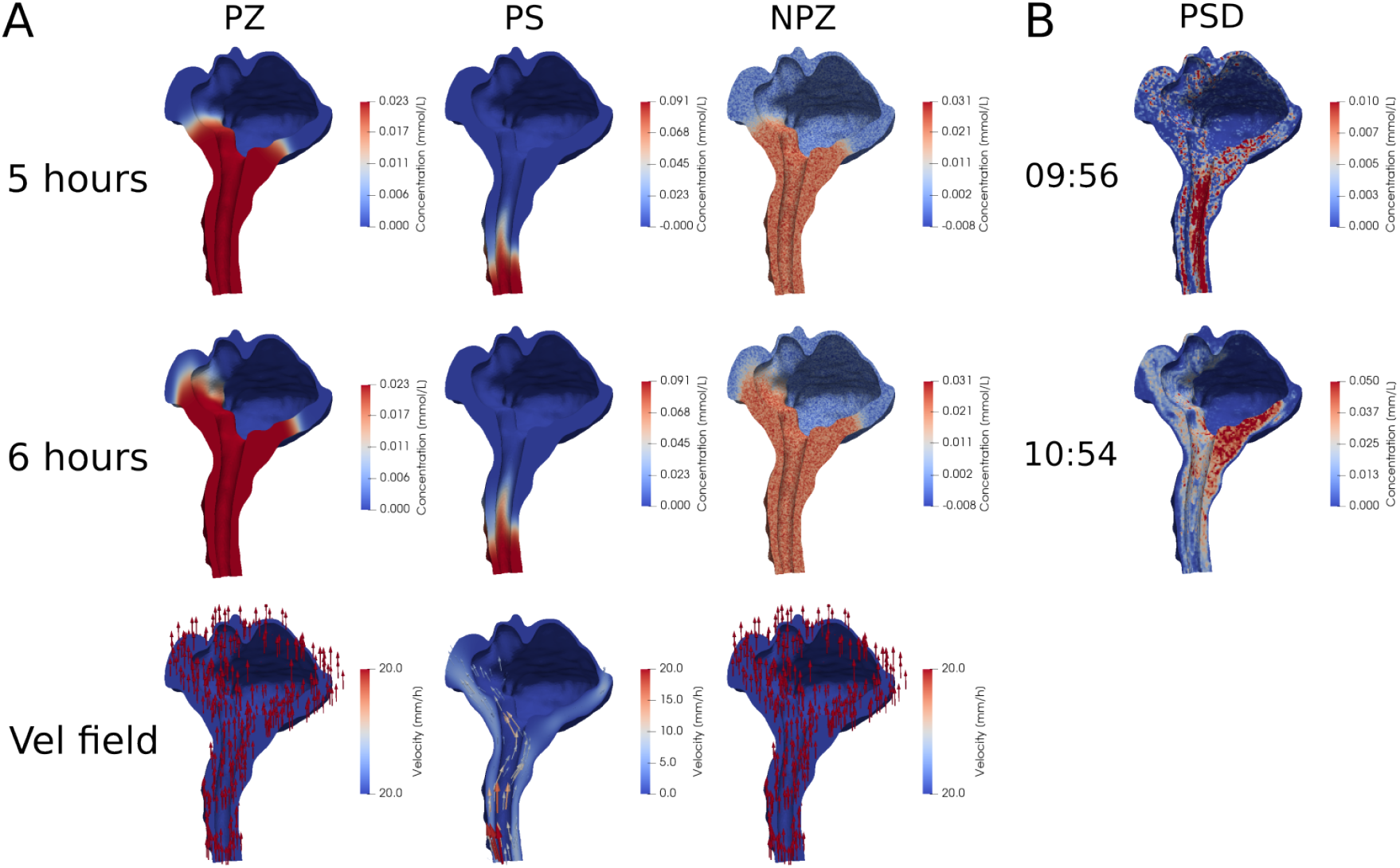
(**A**) The manufactured concentration images for the PZ, PS and NPZ problems shown at two time-points (top two rows) with the corresponding velocity fields used to create them (bottom row). The second image at 6 hours is the longest computed timestep. Note that the velocity field for the PZ and NPZ problem is the same. (**B**) The concentration images used in the PSD problem. The image times are 09:56 and 10:54 o’clock. The scale for the PSD problem is cut at 0.010 and 0.050 for the first and second time-point respectively, in a way such that any point with concentration larger than the cutoff is marked in red. This is to make the figures more readable as the PSD problem has concentration spikes in small areas.

### 2.9 Numerical methods and simulation software

The OCDC methods are implemented using PDE constrained optimization in *dolfin adjoint*^28^. In all cases the concentrations, *c*, and velocity field, *ϕ*, are represented as continuous piecewise linear functions. Time discretization is done through the backwards Euler method. The optimization method is terminated when the norm of the projected gradient is less than 10^−8^ or after 200 iterations.

The forward problem is solved using FEM with the same spaces and discretization as for OCDC and OMT; implemented using the FEniCS finite elements software^29,30^.

Including testing and validation, a total of 15 000 CPU hours were used to run the simulations. All simulations were run on high-performance computing infrastructure Sigma2 - the National Infrastructure for High Performance Computing and Data Storage in Norway.

### 2.10 Concentration estimates

The tracer concentration was estimated following the procedure described in^20^, which used the signal increase and the initial T1 time to map to a given concentration. The T1 time for the CSF was set to 5000 ms, and the relaxivity rate *r*1 was set 3.2 L/mmols^31^. The T1 times for CSF was reported to be over 3000 ms^32^ and close to 4500 ms^33^ at 3T, but difference by using 5000 ms instead of 4500 ms is less than 1 *µ*mol*/*L.

## 3 Results

The character and values of the reconstructed velocity field vary depending on the different methodologies and parameters. In the following section, we focus exclusively on the highest resolution mesh for both the box and patient mesh. The full dataset accounting for all parameters is included in Appendix **??**.

### 3.1 Reconstructed velocity from manufactured problems on a simple box mesh

For each of the four methods, we find a velocity field for the box mesh by solving the manufactured problems. We characterize the velocity field by two values, namely, the maximum and average velocity. For the BZ problem the true maximum and average velocities are both 20 mm/h and for the BS problem the true maximum velocity is 20 mm/h and the true average velocity is 9.50 mm/h. The problems are solved for different resolutions and timesteps between the first and second image (Section 2.8), but this section only presents the results for the highest resolution mesh (26364 cells). This section mainly focuses on the BZ and BS problem, because the NBZ problem is purely intended to study the error introduced by noise on the BZ problem.

For the BZ problem the OCDC and OCDCz methods finds velocities close to the true velocity and the results largely do not depend on the regularization parameter, for the shortest timestep of 5.67 minutes (Table 3). The OCDC method results in a maximum velocity is in the range 19.94-20.85 mm/h and the average velocity is in the range 18.04-20.31 mm/h depending on the regularization parameter. In the same vein, the maximum velocity is in the range 19.85-20.67 mm/h and the average velocity is in the range 17.87-20.01 for the OCDCz method. The OCDCs method shows worse results with the maximum velocity found to be 30.07 mm/h and the average 14.28 mm/h. Last, the OMT method has a larger dependence on the strength of the regularization with the maximum velocity found to be in the range 3.88-23.23 mm/h and the average velocity in the range 0.52 mm/h-4.96 mm/h.

**Table 3.** The maximum, and average velocity, in parenthesis, of the velocity field for the different methods with a range of regularization parameters and the highest resolution mesh. Note that for the OCDCs method the regularization parameter is 0. The timestep between the images is 5.67 minutes. The velocity fields are generated using the methods described in section 2, the true maximum velocity is 20 mm/h for both the BZ and BS problem and the true average velocity is 20 mm/h for the BZ problem and 9.50 mm/h for the BS problem.

| Problem | $\alpha$<br>Method | $10^{-4}$ | $10^{-5}$ | $10^{-6}$ | 0 |
| --- | --- | --- | --- | --- | --- |
| BZ | OCDC | 20.41 (20.31) | 19.94 (19.40) | 20.85 (18.04) | - |
|  | OCDCz | 19.85 (19.79) | 20.51 (20.01) | 20.67 (17.87) | - |
|  | OCDCs | - | - | - | 30.07 (14.28) |
|  | OMT | 3.88 (0.52) | 15.99 (2.59) | 23.23 (4.96) | - |
| BS | OCDC | 10.39 (10.29) | 12.22 (11.40) | 14.61 (11.94) | - |
|  | OCDCz | 10.06 (9.95) | 11.19 (10.26) | 15.03 (10.36) | - |
|  | OCDCs | - | - | - | 20.77 (9.86) |
|  | OMT | 9.81 (0.79) | 19.32 (1.59) | 21.52 (2.26) | - |

The OCDC and OCDCz methods show a slightly larger dependence on the regularization parameters when solving the BS problem (Table 3). For this problem, the OCDC method finds a maximum velocity in the range 10.39-14.61 mm/h and an average velocity in the range 10.29-11.94 mm/h. The OCDCz method finds a maximum velocity in the range 10.06-14.61 mm/h and an average velocity in the range 9.95-15.03 mm/h. The maximum velocity is 20.77 mm/h and the average velocity is 9.86 mm/h for the OCDCs method. This is very close to the true velocity.

Finally, the OMT method again depends on the choice of regularization with a maximum velocity in the range 9.81-21.52 mm/h and the average velocity in the range 0.79-2.26 mm/h. A visualization of selected velocity field solutions are shown in Figure 4.

**Figure 4.**
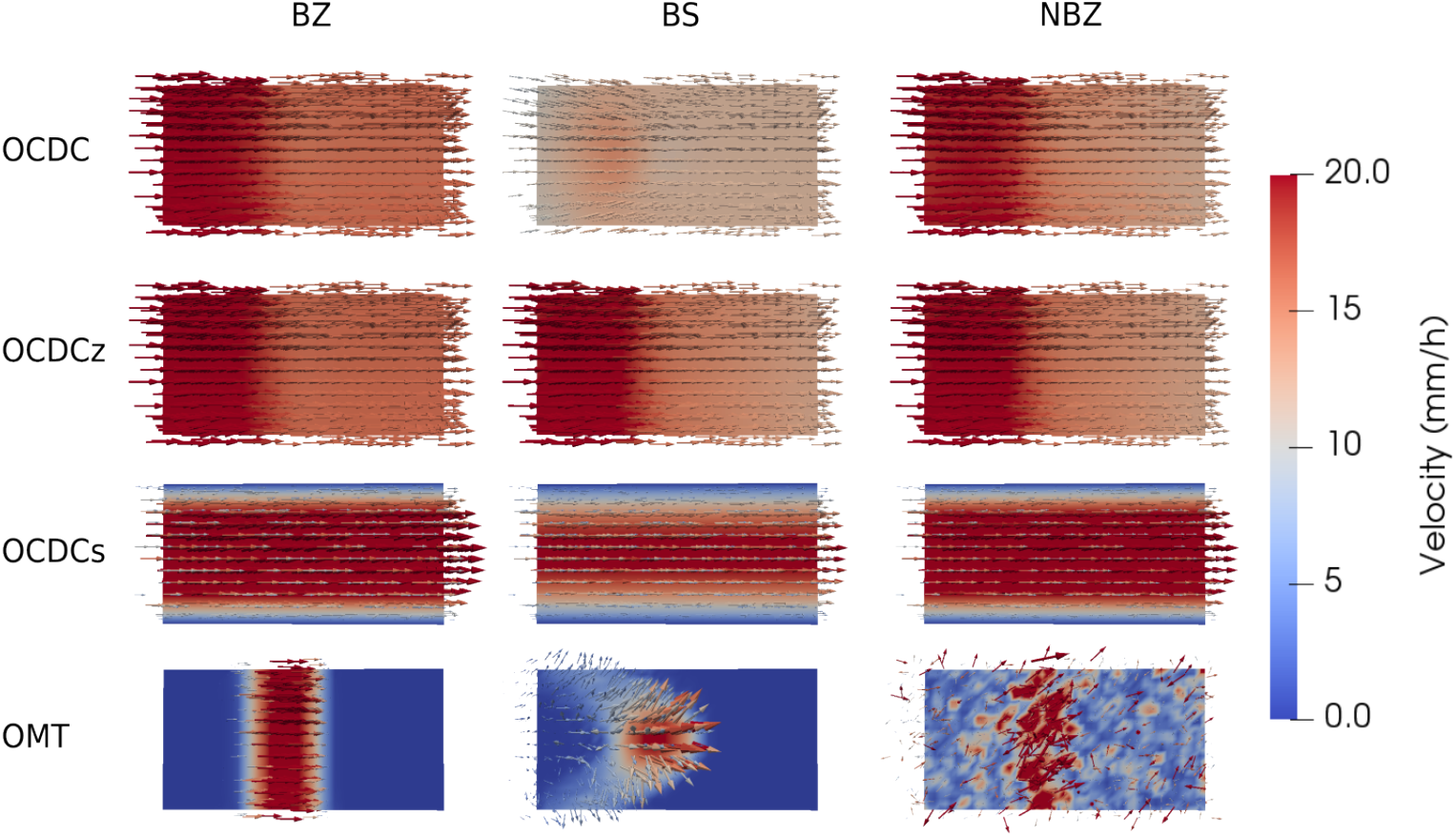
Visual representation of the velocity fields found for the manufactured problems on the box mesh. Each row shows one method and each column one problem. The regularization parameter is 10^−6^, except for the OCDCs method where the regularization parameter is 0. In all cases the timestep is 5.67 minutes. The true velocity field is shown the bottom row of Figure 2.

The velocities closest to the true velocity, with a regularization parameter of 10^−6^, for the BZ problem are found with the OCDC and OCDCz methods with the shortest timestep of 5.67 minutes (Table 4). They find a maximum and average velocity of 20.85 mm/h and 18.04 mm/h, and 20.67 mm/h and 17.87 mm/h respectively. For longer timesteps the average velocity is underestimated while the maximum velocity does not change considerably. The OCDCs method overestimates the maximum velocity for the shortest timestep, finding a maximum velocity of 30.07 mm/h and an average velocity of 14.28 mm/h. For larger timesteps both the maximum and average velocity decrease.

**Table 4.** The maximum, and average velocity, in parenthesis, of the velocity field for the different methods with a range of timesteps (abbreviated as ts in the table) using the highest resolution mesh. The regularization is 10^−6^, except of the OCDCs method which has a regularization of 0. The velocity fields are generated using the methods described in section 2, the true maximum velocity is 20 mm/h for both BZ and BS and the true average velocity is 20 mm/h for the BZ method and 9.50 mm/h for the BS method.

| Problem | Method \ ts (min) | 5.67 | 11.25 | 16.92 | 22.5 |
| --- | --- | --- | --- | --- | --- |
| BZ | OCDC | 20.85 (18.04) | 20.57 (13.90) | 20.29 (12.21) | 20.09 (12.02) |
|  | OCDCz | 20.67 (17.87) | 20.55 (13.92) | 20.28 (12.21) | 20.04 (11.97) |
|  | OCDCs | 30.07 (14.28) | 20.58 (9.77) | 16.16 (7.67) | 13.65 (6.48) |
|  | OMT | 23.23 (4.96) | 34.07 (6.48) | 30.93 (7.44) | 22.10 (6.21) |
| BS | OCDC | 14.61 (11.94) | 13.57 (9.83) | 12.38 (8.17) | 11.52 (7.39) |
|  | OCDCz | 15.03 (10.36) | 17.18 (8.48) | 18.38 (7.07) | 18.98 (6.30) |
|  | OCDCs | 20.77 (9.86) | 18.63 (8.85) | 15.35 (7.29) | 13.22 (6.28) |
|  | OMT | 21.52 (2.26) | 30.04 (2.39) | 46.77 (2.62) | 56.39 (2.87) |

The OMT method varies with the length of the timestep, having a maximum velocity in the range 22.10-34.07 mm/h and average velocity in the range 4.96-7.44 mm/h.

For the BS problem, the best results are found with the shortest timestep (Table 4). Here, the OCDCs method finds the true solution almost perfectly with a maximum velocity of 20.77 mm/h and average velocity of 9.86 mm/h. The OCDC and OCDCz methods underestimate the maximum and average velocity finding a maximum and average velocity of 14.61 mm/h and 11.94 mm/h, and 15.03 mm/h and 10.36 mm/h respectively. While the OMT method finds a quite accurate maximum solution of 21.52 mm/h, but severely underestimates the average velocity at 2.26 mm/h. All the methods find worse solutions for larger timesteps.

#### 3.1.1 Method convergence and solution dependency on noise

In order to judge how successful the different OCDC methods are at recreating the velocity fields, we look at the reduction in the *J*-functional (9) over 200 iterations. The *J*-functional reduction for each method on each problem will depend on the choice of regularization parameter, mesh resolution and timestep. For the BZ problem the *J*-functional reduction is at least larger than 91.54 % for the OCDC method, 91.66 % for the OCDCz method and

69.04 % for the OCDCs method (Table 5). Similarly, the *J*-functional reduction is larger than 74.00 % for the OCDC method, 72.29 % for the OCDCz method and 95.75 % for the OCDCs method for the BS problem (Table 5). Note that the OMT method does not have a *J*-functional associated with it and is therefore not assessed in this way.

**Table 5.** The Reduction of *J*, (9), for the different OCDC methods with a range of regularization parameters using the highest resolution mesh. The time difference between the images is 5.67 minutes.

| Problem | Method \ $\alpha$ | $10^{-4}$ | $10^{-5}$ | $10^{-6}$ | 0 |
| --- | --- | --- | --- | --- | --- |
| BZ | OCDC | 91.54 | 92.12 | 94.74 | - |
|  | OCDCz | 91.66 | 92.08 | 94.75 | - |
|  | OCDCs | - | - | - | 69.04 |
| BS | OCDC | 74.00 | 82.68 | 94.37 | - |
|  | OCDCz | 72.29 | 75.41 | 85.40 | - |
|  | OCDCs | - | - | - | 95.75 |

To investigate the methods robustness with respect to noise, we solve the NBZ problem and look at the *L*_2_ difference, described in Section 2.7, to the solution the BZ problem. This is done for all regularization parameters and timesteps. For the average and maximum velocity the maximum *L*_2_ difference in solution is 7.3 % and 7.9 % for the OCDC method, 15.9 % and 8.6 % for the OCDCz method, 95.1 % and 89.8 % for the OCDCs method and 620 % and 330 % for the OMT method.

### 3.2 Velocity field reconstructed from manufactured problems on a patient mesh

We tested the methods on a realistic geometry by solving the PZ and PS problems (Section 2.3). As for the box mesh, we characterize the velocity fields by the maximum and average velocity. For the PZ problem, the true maximum and average velocities are both 20 mm/h and for the PS problem the true maximum and average velocities are 20 mm/h and 2.77 mm/h respectively. The problems are solved for different resolutions and timesteps between the two images, but this section, as was the case for the former, focuses on the highest resolution. This section also focuses mostly on the PZ and PS problem. Note that the regularization parameters are 100 times larger for the patient mesh compared to the box mesh. This choice was made because the patient mesh has a 50.3 times larger volume and was confirmed to be a good choice based on numerical experimentation where a range of parameters was tested to fnd a point of stability.

The solutions of the PZ problem are more dependent on the regularization parameter compared to the BZ problem (Section 3.1) for the shortest timestep; here 15 minutes (Table 6). Compared to the corresponding problem on the box mesh we also note that the average velocity is always underestimated. For our regularization parameters, the OCDC method finds a maximum velocity in the range 13.09-29.80 mm/h and an average velocity in the range 4.56-8.19 mm/h. OCDCz, similarly finds a maximum velocity in the range 4.05 mm/h-18.26 mm/h and an average velocity in the range 0.45-0.92 mm/h, and OCDCs finds 54.06 mm/h and 7.49 mm/h as the maximum and average velocity respectively. Lastly, OMT finds a maximum velocity in the range 13.89-33.30 mm/h and an average velocity in the range 0.68-1.54 mm/h.

**Table 6.** The maximum, and average velocity, in parenthesis, of the velocity field for the different methods with a range of regularization parameters using the highest resolution mesh. The time difference between the images is 15 minutes. The velocity field is generated using the methods described in section 2, the true maximum velocity is 20 mm/h for both PZ and PS and the true average velocity is 20 mm/h for the PZ method and 2.77 mm/h for the PS method.

| Problem | $\alpha$<br>Method | $10^{-6}$ | $10^{-7}$ | $10^{-8}$ | 0 |
| --- | --- | --- | --- | --- | --- |
| PZ | OCDC | 13.09 (4.56) | 25.52 (7.77) | 29.80 (8.19) | - |
|  | OCDCz | 4.05 (0.45) | 8.94 (0.87) | 18.26 (0.92) | - |
|  | OCDCs | - | - | - | 54.06 (7.49) |
|  | OMT | 13.89 (0.68) | 20.92 (1.13) | 33.30 (1.54) | - |
| PS | OCDC | 8.46 (1.61) | 9.86 (1.34) | 11.64 (1.24) | - |
|  | OCDCz | 13.40 (0.39) | 33.45 (0.47) | 74.64 (0.54) | - |
|  | OCDCs | - | - | - | 20.65 (2.86) |
|  | OMT | 8.85 (0.33) | 12.93 (0.43) | 29.35 (0.53) | - |

Similarly, the PS problem solutions are more dependent on the regularization parameter (Table 6) compared to its box counterpart. The OCDC method finds a maximum velocity in the range 8.46-11.64 mm/h, and an average velocity in the range 1.24-1.61 mm/h. Further, the OCDCz method finds maximum a velocity in the range 13.40-74.64 mm/h and an average velocity in the range 0.39-0.54 mm/h, the OCDCs method finds a maximum velocity of 20.65 mm/h and an average velocity of 2.86 mm/h and finally, the OMT method finds a maximum velocity in the range 8.85-29.35 mm/h and an average velocity in the range 0.33-0.53 mm/h. The velocity fields reconstructed with all methods on these two manufactured problems are displayed in the first two columns of Figure 6.

For a regularization parameter of 10^−7^, the solutions of the PZ problem are relatively stable with respect to timestep length (Table 7), but less so than for the box mesh (Section 3.1). The OCDC and OMT method finds their best results for the shortest timestep, that is a maximum and average velocity of 25.52 mm/h and 7.77 mm/h, and 20.92 mm/h and 1.13 mm/h respectively. The solution found using the OCDCz method increase more with increasing timestep finding maximum velocities in the range 8.94-24.28 mm/h and average velocities in the range 0.70-0.87 mm/h. The OCDCs method overestimates the maximum velocity for all timestep lengths finding at best a maximum velocity of 33.99 mm/h and an average velocity of 4.71 mm/h for the longest timestep.

**Table 7.** The maximum, and average velocity, in parenthesis, of the velocity field for the different methods with a range of timesteps (abbreviated as ts in the table) and the highest resolution mesh. The regularization is 10^−7^, except of the OCDCs method which has a regularization of 0. The velocity field is generated using the methods described in section 2. The true maximum velocity is 20 mm/h for both PZ and PS and the true average velocity is 20 mm/h for the PZ method and 2.77 mm/h for the PS method.

| Problem | ts (min)<br>Method | 15 | 30 | 45 | 60 |
| --- | --- | --- | --- | --- | --- |
| PZ | OCDC | 25.52 (7.77) | 27.41 (8.30) | 28.06 (8.65) | 28.21 (8.89) |
|  | OCDCz | 8.94 (0.87) | 14.75 (0.76) | 19.17 (0.70) | 24.28 (0.70) |
|  | OCDCs | 54.06 (7.49) | 41.86 (5.80) | 37.75 (5.23) | 33.99 (4.71) |
|  | OMT | 20.92 (1.13) | 24.18 (1.19) | 31.34 (1.32) | 32.90 (1.50) |
| PS | OCDC | 9.86 (1.34) | 10.52 (1.29) | 10.32 (1.27) | 10.22 (1.26) |
|  | OCDCz | 33.45 (0.47) | 54.81 (0.61) | 64.88 (0.68) | 70.83 (0.73) |
|  | OCDCs | 20.31 (2.81) | 20.51 (2.84) | 20.65 (2.86) | 20.75 (2.88) |
|  | OMT | 12.93 (0.43) | 13.11 (0.43) | 13.25 (0.44) | 13.61 (0.44) |

With a regularization parameter of 10^−7^ the solutions of the PS problem are more independent of the timestep length than for the PZ problem. The OCDC and OMT method both underestimates the maximum and average velocity, finding a maximum velocity in the range 9.86-10.22 mm/h and an average velocity in the range 1.26-1.34 mm/h, and a maximum velocity in the range 12.93-13.61 mm/h and an average velocity in the range 0.43-0.44 mm/h respectively. The OCDCz method overestimates the velocity in all cases, but finds the best solution of a maximum velocity of 33.45 mm/h and an average velocity of 0.47 mm/h for the smallest timestep. The OCDCs is very close to the actual solution for all timesteps and finds the best solution for the smallest timestep which is a maximum velocity of 20.31 mm/h and an average velocity of 2.81 mm/h.

Note that as opposed to the box mesh the maximum and average velocity generally increase with timestep length, except for the OCDCs method on the PZ problem.

#### 3.2.1 Method convergence and solution dependency on parameters and noise

The convergence of the solutions is assessed by the reduction in the *J*-functional (9) over 200 iterations. For the PZ problem the three methods OCDC, OCDCz and OCDCs have a minimum reduction of 74.70 %, 9.89 %, 60.19 % and for the PS they have a minimum reduction of 97.04 %, 46.41 % and 99.71 % respectively (Table 8), for the velocities presented in (Table 6). Note, as earlier, that the OMT method does not have a *J*-functional associated with it and is therefore not assessed this way.

**Table 8.** The Reduction of the *J*-functional, defined in (9), for the different OCDC methods with a range of regularization parameters using the highest resolution mesh.

| Problem | $\alpha$<br>Method | $10^{-6}$ | $10^{-7}$ | $10^{-8}$ | 0 |
| --- | --- | --- | --- | --- | --- |
| PZ | OCDC | 74.70 | 94.98 | 99.22 | - |
|  | OCDCz | 9.89 | 12.51 | 13.77 | - |
|  | OCDCs | - | - | - | 60.19 |
| PS | OCDC | 97.04 | 99.19 | 99.75 | - |
|  | OCDCz | 46.41 | 59.66 | 66.78 | - |
|  | OCDCs | - | - | - | 99.71 |

The methods resistance to noise is measured by comparing the *L*_2_ difference as described in Section 2.7 for the maximum and average velocity in the NPZ and PZ problems for the four methods. The maximum *L*_2_ difference found for the maximum and average velocity is 13.5 % and 18.3 % for the OCDC method, 279.1 % and 52.2 % for the OCDCz method, 24.6 % and 24.5 % for the OCDCs method, and 755 % and 173 % for the OMT method.

The dependence of the PZ problem on regularization is shown in (Figure 5). We remark that the reconstructed velocity field becomes more unstable for smaller regularization parameters (not shown in the figure).

**Figure 5.**
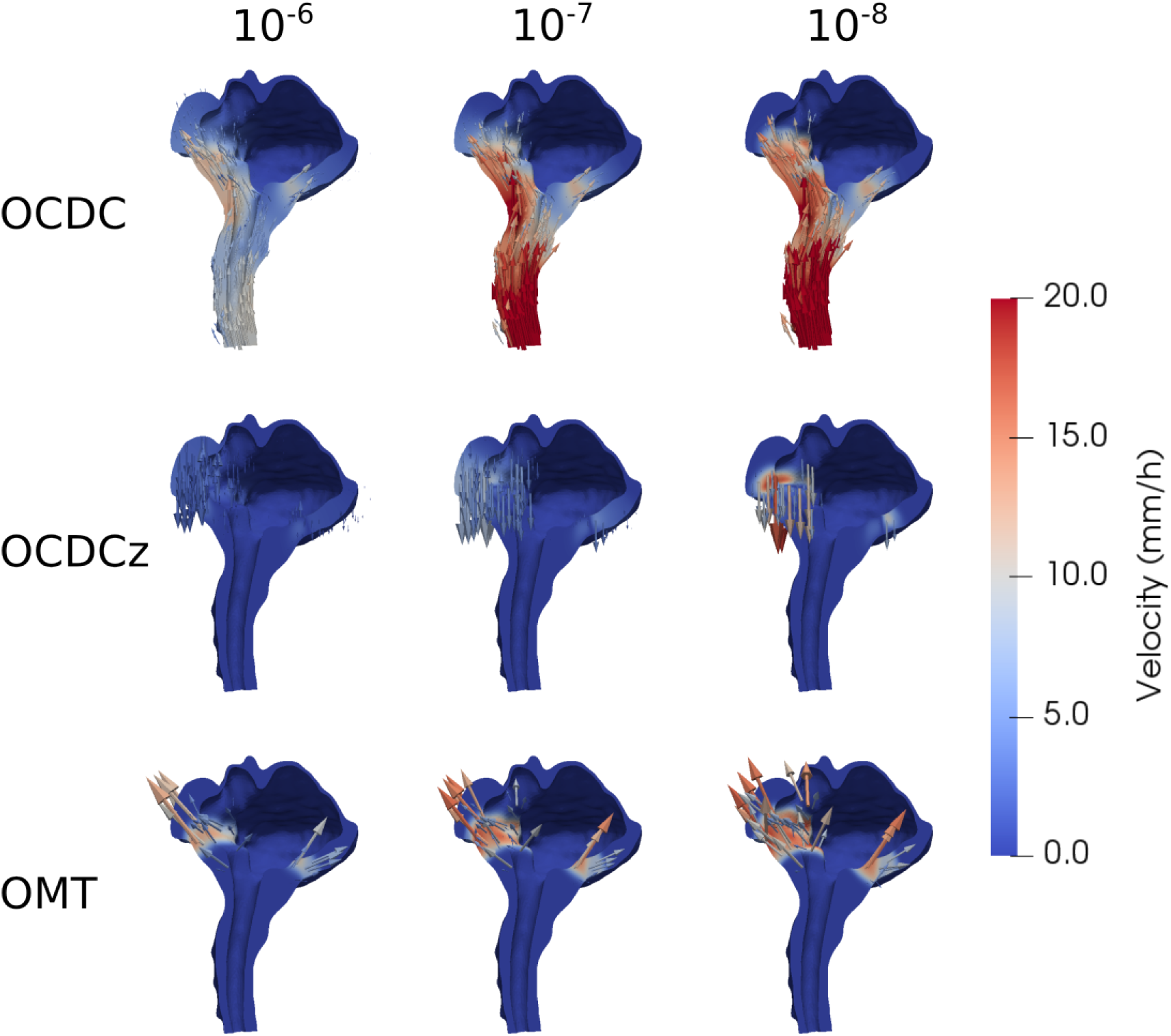
The reconstructed velocity fields for the PZ problem with the OCDC, OCDCz and OMT methods for different regularization parameters. The highest resolution mesh is used and the timestep is 15 minutes for the manufactured problems. Note that the velocity scale is cut off at 20 mm/h, such that any velocity above will appear in red. The true velocity field for PZ is shown in Figure 3.

### 3.3 Velocity field reconstructed from patient data on a patient mesh

For each of the four methods, we find a velocity field for the PSD problem. The reconstructed velocity field depends on the regularization parameter and the resolution of the mesh, but we focus here exclusively on the largest mesh. As earlier, we quantify the fields by the maximum and average velocity.

The solutions depends more heavily on the regularization parameters than for the manufactured problems (Table 9). For the OCDC method, the maximum velocity is in the range 14.75-46.92 mm/h and average velocity is in the range 1.89-2.41 mm/h. For the OCDCz method the maximum velocity is in the range 25.39-79.70 mm/h and the average velocity is in the range 0.08-0.11 mm/h. The maximum velocity is 79.06 mm/h and the average velocity is 10.96 mm/h for the OCDCs method and, finally, the maximum velocity is in the range 9.22-54.34 mm/h and the average velocity is in the range 0.049-1.25 mm/h for the OMT method. The velocity fields found by all methods are shown in the last column of Figure 6.

**Figure 6.**
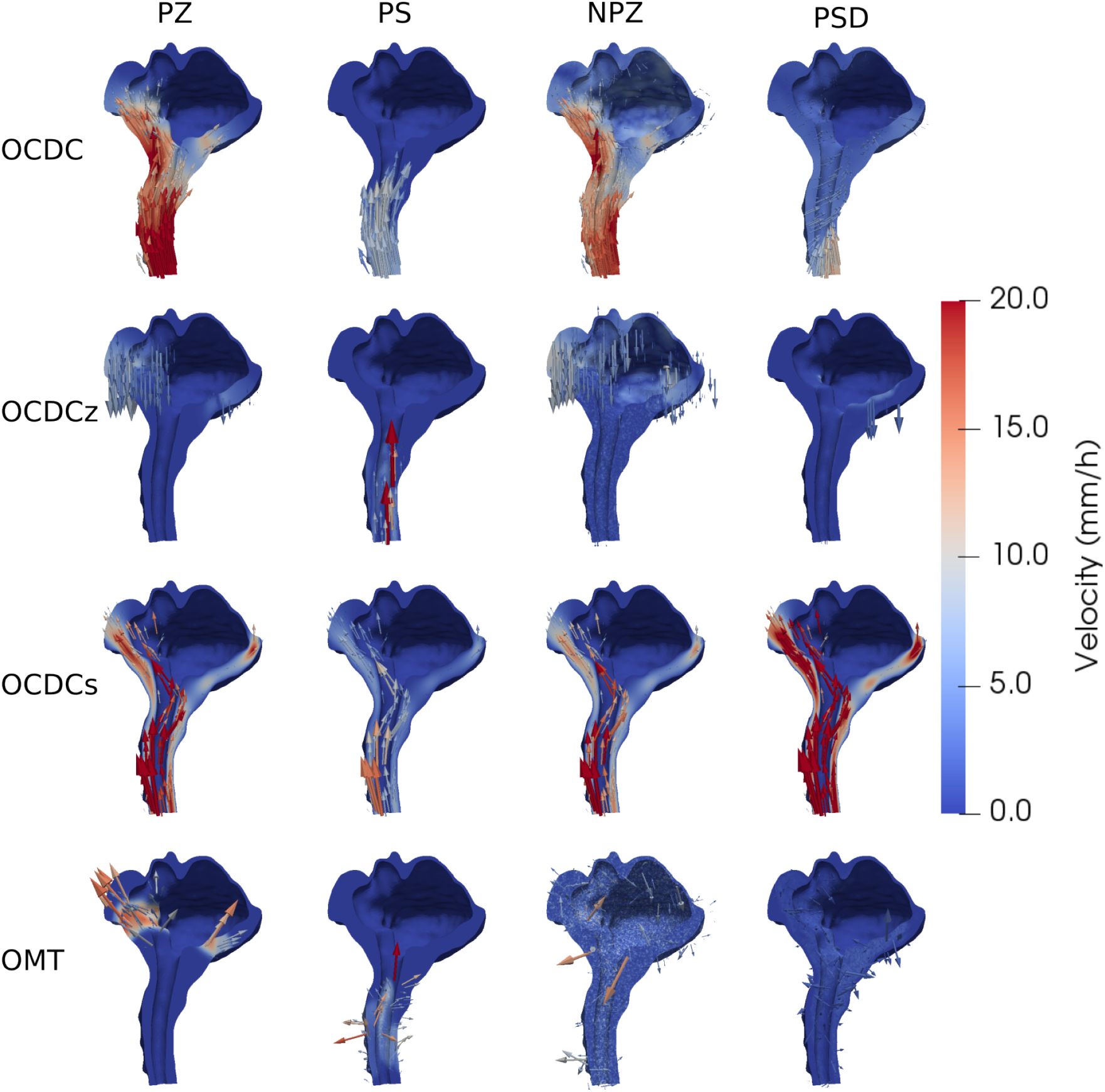
Visual representation of the velocity fields found for the PSD and manufactured problems. Each row shows one method and each column one problem. For the PSD problem the regularization parameter is 10^−4^ in all cases and for the manufactured problem columns the regularization parameter is 10^−7^, except for the OCDCs method where the regularization parameter is 0. The highest resolution mesh is used and the timestep is 15 minutes for the manufactured problems. Note that the velocity scale is cut off at 20 mm/h, such that any velocity above will appear in red. The true velocity field for PZ, PS NPZ is shown in Figure 3.

**Table 9.** The maximum, and average velocity, in parenthesis, of the velocity field for the different methods with a range of regularization parameters and the highest resolution mesh. The timestep between the images are 57 minutes and 48 seconds. The velocity fields are generated using the methods described in section 2 on patient data.

| Problem | $\alpha$<br>Method | $10^{-4}$ | $10^{-5}$ | $10^{-6}$ | 0 |
| --- | --- | --- | --- | --- | --- |
| PSD | OCDC | 14.75 (2.41) | 23.96 (2.14) | 46.92 (1.89) | - |
|  | OCDCz | 25.39 (0.08) | 29.91 (0.08) | 79.70 (0.11) | - |
|  | OCDCs | - | - | - | 79.06 (10.95) |
|  | OMT | 9.22 (0.49) | 19.23 (0.97) | 54.34 (1.25) | - |

#### 3.3.1 Method convergence and solution dependency on parameters

To asses whether our methods are converging, we look at the reduction of the *J*-functional (9) over 200 iterations. The OCDC and OCDCz methods have a *J*-functional reduction of up between 19.47 % and 28.49 %, and 13.64 % and 22.47 % respectively, while the OCDCs method only has a reduction of 3.44 % at max (Table 10). This is in all cases a considerably smaller *J*-functional reduction than for the manufactured solutions.

**Table 10.** The Reduction of the *J*-funcional, (9), for the different OCDC methods with a range of and regularization parameters using the highest resolution mesh. Note that for the OCDCs method the regularization parameter is 0.

| Method \ $\alpha$ | $10^{-4}$ | $10^{-5}$ | $10^{-6}$ | 0 |
| --- | --- | --- | --- | --- |
| OCDC | 19.47 | 25.53 | 28.49 | - |
| OCDCz | 13.64 | 18.87 | 22.47 | - |
| OCDCs | - | - | - | 3.44 |

The velocity fields change significantly with change in the regularization parameter (Figure 7). By visual inspection, we can see that as the regularization parameter goes toward 10^−6^ the velocity field becomes increasingly irregular and there occurs rapid spacial changes in velocity. For the OMT method, specifically, the solution is irregular for all regularization parameters, but there is still a significant increase in rapid velocity changes for small regularization parameters.

**Figure 7.**
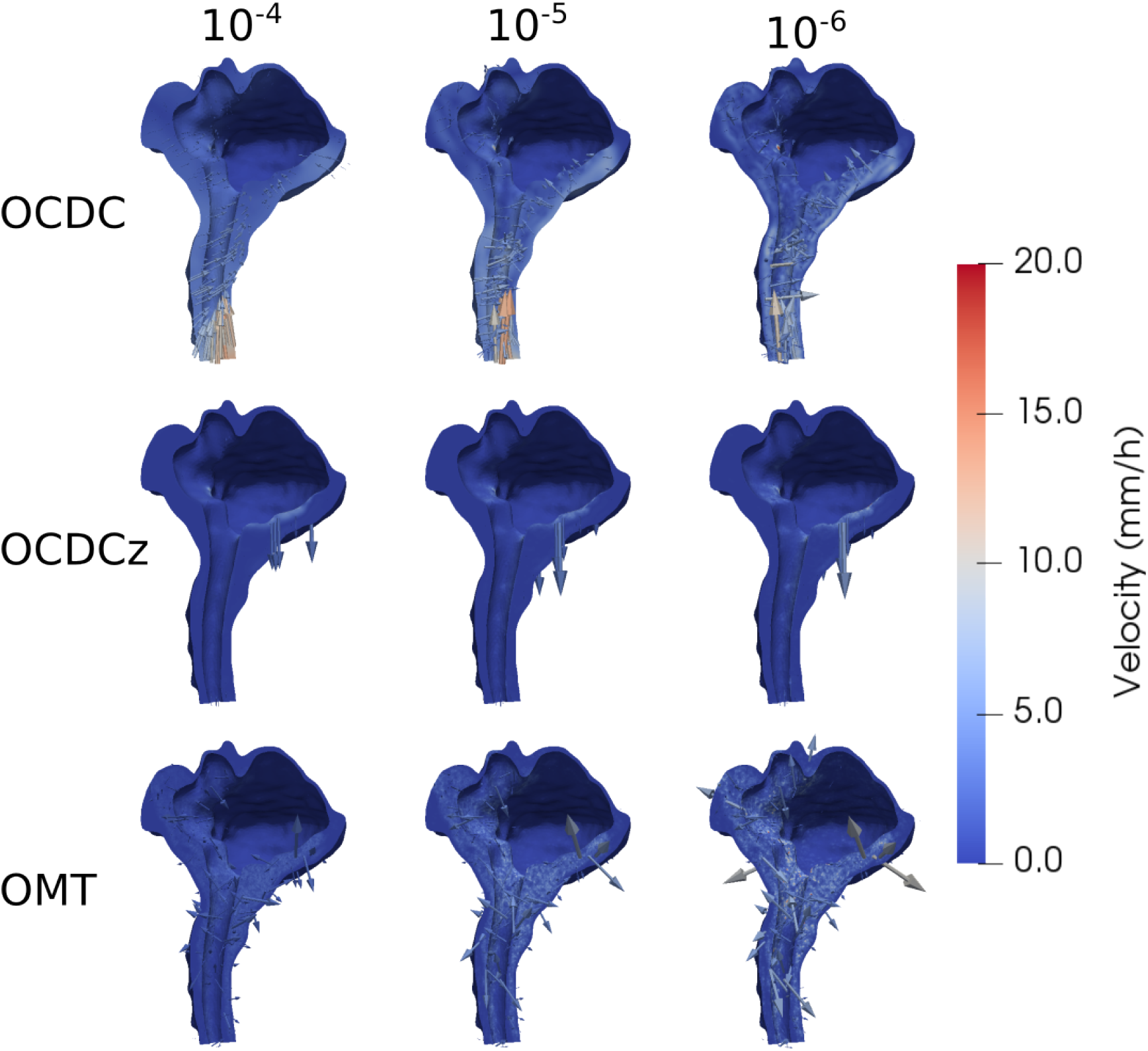
The reconstructed velocity fields for the PSD problems with the OCDC, OCDCz and OMT methods for different regularization parameters. Each row shows a method and each column shows a regularization parameter. The highest resolution is used. Note that the velocity scale is cut off at 20 mm/h, such that any velocity above will appear in red.

## 4 Discussion

In this paper we have used several versions of an OCDC method and an OMT method to recover velocity fields from images. We have explored images based on manufactured problems as well as patient data. The manufactured problems are posed on a box mesh and on a realistic lower brain mesh. For the box mesh, the solutions are relatively independent of the regularization parameter, in the range of 10^−4^ − 10^−6^ for a timestep of 5.67 minutes. The OMT method is a bit of an exception, with the maximum velocity increasing almost sixfold with the regularization parameter. In general, although this varies across methods and problems, a good solution is found with a regularization parameter of 10^−6^. For this regularization parameter the solutions are also stable with respect to timstep length, with the OCDCs method being a bit of an exception. The solutions are generally better for shorter timestep lengths.

The solutions show more variance with regularization parameter for the manufactured solutions on the patient mesh than for the box mesh. In the regularization parameter range 10^−6^ − 10^−8^ the methods find maximum velocities close to the actual solution, but the average velocity is in general underestimated. Solutions that are stable with respect to timestep length are found for a regularization parameter of 10^−7^, with some exceptions. The velocity fields found by the methods look reasonable in some regions, but in regions with low or zero concentration change between images the velocity is close to zero, explaining the underestimation of the average velocity (Figure 6). The patient mesh is roughly 50 times larger, in volume, compared to the box mesh which is probably the reason we experience this problem here and not in the box mesh.

Note that, though the reconstructed velocity fields are inaccurate, the maximum and average velocities are in general of the correct order of magnitude. That is, within a factor of 10 of the correct solution.

We similarly observe that the reconstructed velocity field is larger in regions with change in concentration from image 1 to 2 when solving the PSD problem. The reduction in the *J*-functional is lower than for the manufactured solution (Table 10), but is slightly better for a small regularization parameter of 10^−6^. However, for this regularization parameter the solution looks very chaotic (Figure 7). Interestingly, we observe higher velocity around the cisterna magna in the mesh which could be explained by this being a possible outflow route^6^.

The maximum velocity found for the PSD problem is between 14.75–79.70 mm/h. At the high end, this is comparable to velocities found by modeling, which has found peak bulk flow velocities between 20 and 50 *µ*m/s in the SAS of humans^11,14^ and is also similar to those found in rodents^2,34^. It is, however, significantly lower than the pulsatile velocities of 5 cm/s found in the foramen magnum. This is to be expected, as the timestep between images is roughly 1 hour and should therefore not capture pulsatile effects which occur on a shorter time-scale.

The OMT method has been used successfully to calculate interstitial fluid transport in rat brains^25,26^. Here, we have more trouble using the OMT and similar methods. A key difference could be the size difference between the rat and human brain. This matches well with our experience on the smaller box mesh where our methods were much more accurate.

## 5 Conclusion

In conclusion, we find that our methods reconstruct velocities well for manufactured problems on a box mesh and we find regions where the solutions are stable in terms of regularization parameter and timestep choice. The methods struggle more with a realistic patient mesh, where they are more dependent on regularization parameter, but for a good choice of regularization parameter the solutions are stable with respect to timestep length. The methods tend to underestimate the average velocity with a very small velocity in regions without concentration change between the two images used in the calculations. On realistic data, the methods fail to reduce the *J*-functional unless the regularization parameter is small, and for these parameters the solution looks chaotic. However, the values of the maximum velocity produced are in line with what we would expect from previous studies. We find therefore that the methods chosen, in their current form, are not satisfactory in solving the biophysics problems presented here and further work needs to explore improvement in the methods.

## Achknowledgements

Special thanks to Marie E. Rognes for developing the mathematics behind the OCDC method and letting us use her python scripts as a starting point for our numerical calculations.

## References

1. Iliff, J. J. et al. A paravascular pathway facilitates CSF flow through the brain parenchyma and the clearance of interstitial solutes, including amyloid β. Sci. translational medicine 4, 147ra111–147ra111 (2012).

2. Mestre, H. et al. Flow of cerebrospinal fluid is driven by arterial pulsations and is reduced in hypertension. Nat. communications 9, 4878 (2018).

3. Bojarskaite, L. et al. Sleep cycle-dependent vascular dynamics in male mice and the predicted effects on perivascular cerebrospinal fluid flow and solute transport. Nat. communications 14, 953 (2023).

4. Ray, L. A., Pike, M., Simon, M., Iliff, J. J. & Heys, J. J. Quantitative analysis of macroscopic solute transport in the murine brain. Fluids Barriers CNS 18, 1–19 (2021).

5. Haga, P. T. et al. A numerical investigation of intrathecal isobaric drug dispersion within the cervical subarachnoid space. PLoS One 12, e0173680 (2017).

6. Eide, P. K., Valnes, L. M., Lindstrøm, E. K., Mardal, K.-A. & Ringstad, G. Direction and magnitude of cerebrospinal fluid flow vary substantially across central nervous system diseases. Fluids Barriers CNS 18, 1–18 (2021).

7. Balédent, O. Imaging of the cerebrospinal. Adult hydrocephalus 256, 121 (2014).

8. Dreha-Kulaczewski, S. et al. Inspiration is the major regulator of human csf flow. J. neuroscience 35, 2485–2491 (2015).

9. Liu, P. et al. Cardiac and respiratory activities induce temporal changes in cerebral blood volume, balanced by a mirror csf volume displacement in the spinal canal. NeuroImage 305, 120988 (2025).

10. Lindstrøm, E. K., Ringstad, G., Mardal, K.-A. & Eide, P. K. Cerebrospinal fluid volumetric net flow rate and direction in idiopathic normal pressure hydrocephalus. NeuroImage: Clin. 20, 731–741 (2018).

11. Vinje, V. et al. Respiratory influence on cerebrospinal fluid flow–a computational study based on long-term intracranial pressure measurements. Sci. reports 9, 9732 (2019).

12. Eide, P. K., Vinje, V., Pripp, A. H., Mardal, K.-A. & Ringstad, G. Sleep deprivation impairs molecular clearance from the human brain. Brain 144, 863–874 (2021).

13. Fultz, N. E. et al. Coupled electrophysiological, hemodynamic, and cerebrospinal fluid oscillations in human sleep. Science 366, 628–631 (2019).

14. Hornkjøl, M. et al. CSF circulation and dispersion yield rapid clearance from intracranial compartments. Front. Bioeng. Biotechnol. 10, 932469 (2022).

15. Boster, K. A. et al. Artificial intelligence velocimetry reveals in vivo flow rates, pressure gradients, and shear stresses in murine perivascular flows. Proc. Natl. Acad. Sci. 120, e2217744120 (2023).

16. Koundal, S. et al. Optimal mass transport with lagrangian workflow reveals advective and diffusion driven solute transport in the glymphatic system. Sci. reports 10, 1990 (2020).

17. Benveniste, H. et al. Glymphatic cerebrospinal fluid and solute transport quantified by MRI and PET imaging. Neuroscience 474, 63–79 (2021).

18. Chen, X., Benveniste, H. & Tannenbaum, A. R. Unbalanced regularized optimal mass transport with applications to fluid flows in the brain. Sci. Reports 14, 1111 (2024).

19. Valnes, L. M. et al. Apparent diffusion coefficient estimates based on 24 hours tracer movement support glymphatic transport in human cerebral cortex. Sci. reports 10, 1–12 (2020).

20. Vinje, V. et al. Human brain solute transport quantified by glymphatic MRI-informed biophysics during sleep and sleep deprivation. bioRxiv 2023–01 (2023).

21. Ringstad, G., Vatnehol, S. A. S. & Eide, P. K. Glymphatic MRI in idiopathic normal pressure hydrocephalus. Brain 140, 2691–2705 (2017).

22. Ringstad, G. et al. Brain-wide glymphatic enhancement and clearance in humans assessed with MRI. JCI insight 3 (2018).

23. Kolesov, I., Karasev, P., Tannenbaum, A. & Haber, E. Fire and smoke detection in video with optimal mass transport based optical flow and neural networks. In 2010 IEEE International Conference on Image Processing, 761–764 (IEEE, 2010).

24. Mueller, M., Karasev, P., Kolesov, I. & Tannenbaum, A. Optical flow estimation for flame detection in videos. IEEE Transactions on image processing 22, 2786–2797 (2013).

25. Ratner, V. et al. Optimal-mass-transfer-based estimation of glymphatic transport in living brain. In Medical Imaging 2015: Image Processing, vol. 9413, 406–411 (SPIE, 2015).

26. Ratner, V. et al. Cerebrospinal and interstitial fluid transport via the glymphatic pathway modeled by optimal mass transport. Neuroimage 152, 530–537 (2017).

27. Rognes, M. E. Mathematical modeling of the human brain (vol II): from glymphatics to deep learning, chap. An introduction to identifying velocity fields from contrast imaging via PDE-constrained optimization (Springer, in review, 2024).

28. Mitusch, S., Funke, S. & Dokken, J. dolfin-adjoint 2018.1: automated adjoints for FEniCS and Firedrake. J. Open Source Softw. 4, 1292 (2019).

29. Logg, A., Mardal, K.-A. & Wells, G. Automated solution of differential equations by the finite element method: The FEniCS book, vol. 84 (Springer Science & Business Media, 2012).

30. Alnæs, M. et al. The FEniCS project version 1.5. Arch. Numer. Softw. 3 (2015).

31. Rohrer, M., Bauer, H., Mintorovitch, J., Requardt, M. & Weinmann, H.-J. Comparison of magnetic properties of MRI contrast media solutions at different magnetic field strengths. Investig. radiology 40, 715–724 (2005).

32. Condon, B. et al. MR relaxation times of cerebrospinal fluid. J. computer assisted tomography 11, 203–207 (1987).

33. Chen, L., Bernstein, M., Huston, J. & Fain, S. Measurements of T1 relaxation times at 3.0 T: implications for clinical MRA. In Proceedings of the 9th Annual Meeting of ISMRM, Glasgow, Scotland, vol. 1391 (2001).

34. Bedussi, B., Almasian, M., de Vos, J., VanBavel, E. & Bakker, E. N. Paravascular spaces at the brain surface: Low resistance pathways for cerebrospinal fluid flow. J. Cereb. Blood Flow & Metab. 38, 719–726 (2018).

